# Physics Augmented U-Net: A High-Frequency Aware Generative Prior for Microscopy

**DOI:** 10.1101/2021.12.01.470743

**Authors:** Jathurshan Pradeepkumar, Mithunjha Anandakumar, Vinith Kugathasan, Andrew Seeber, Dushan N. Wadduwage

**Author notes:** These authors contributed equally to the work (Jathurshan Pradeepkumar, Mithunjha Anandakumar, Vinith Kugathasan).

## Abstract

A key challenge in optical microscopy is to image fast at high-resolution. To address this problem, we propose “Physics Augmented U-Net”, which combines deep learning and structured illumination microscopy (SIM). In SIM, the structured illumination aliases out-of-band high-frequencies to the passband of the microscope; thus SIM captures some high-frequencies even when the image is sampled at low-resolution. To utilize these features, we propose a three-element method: 1) a modified U-Net model, 2) a physics-based forward model of SIM 3) an inference algorithm combining the two models. The modified U-Net architecture is similar to the seminal work, but the bottleneck is modified by concatenating two latent vectors, one encoding low-frequencies (LFLV), and the other encoding high-frequencies (HFLV). LFLV is learned by U-Net contracting path, and HFLV is learned by a second encoding path. In the inference mode, the high-frequency encoder is removed; HFLV is then optimized to fit the measured microscopy images to the output of the forward model for the generated image by the U-Net. We validated our method on two different datasets under different experimental conditions. Since a latent vector is optimized instead of a 2D image, the inference mode is less computationally complex. The proposed model is also more stable compared to other generative prior-based methods. Finally, as the forward model is independent of the U-Net, Physics Augmented U-Net can enhance resolution on any variation of SIM without further retraining.

## 1. Introduction

High-resolution fluorescence microscopy is an essential tool in today’s biology. Many important biological molecules can be fluorescently labelled, and their spatial and temporal behavior can be studied. For instance, high-resolution fluorescence imaging can visualize DNA damage and repair process in live cells at sub-micrometer resolution. Moreover, genetically modified animals expressing fluorescence allow molecular specific in-vivo studies, such as the dynamic signaling of neurons in the mouse brain. For many studies requiring high-resolution, rapid image acquisition is fundamentally important. A recent study imaged a cleared whole mouse brain at 6.5*μm*^3^ resolution in 7 minutes [28]. Same imaging at high, diffraction-limited resolution would take over two months. Therefore, the resolution is often sacrificed in favor of scale and time or vice versa. A fluorescence microscopy technique which allows both high-resolution and high-speed has been a vital aim for many decades.

Structured Illumination Microscopy (SIM) is one of the most important high-resolution microscopy techniques developed to date. In contrast to confocal techniques, SIM allows rapid 3D wide-field imaging fully utilizing state-of-the-art cameras. In SIM, multiple structured illumination patterns encode spatial frequencies of the specimen being imaged. The set of encoded images constitute a linear system; by solving the linear system, a depth-resolved image can be reconstructed at high-resolution. Nevertheless, to perform the reconstruction using such traditional solvers, illumination structures should be resolved on the camera according to the Nyquist criterion. When the Nyquist criterion is broken, out-of-band high-frequencies aliases to the passband of the microscope rendering the linear system unsolvable. However, recent advances in deep learning allow solving such under-determined systems by learning content-dependent image priors.

The advancement of deep learning benefited a variety of disciplines, including fluorescence microscopy. In particular, deep convolutional neural networks (DCNNs) can enhance image resolution. Generally, DCNNs learn a mapping from low-resolution images to their respective high-resolution images. This by itself is solving an underdetermined system, but with the absence of any high-frequency information. Undersampled SIM images, however, contain aliased high-frequency information. Thus, DCNNs may translate low-resolution (i.e. undersampled) SIM images to high-resolution images, better than they can translate normal low-resolution images to high-resolution. A DCNN can be trained to translate low-resolution SIM images to high-resolution by acquiring paired training images. However, for each SIM system, such data need to be collected. On the other hand, a mathematical model of the imaging system can be used to create synthetic training data from only a high-resolution image dataset when the model of the system is known a priori. Such models are called forward models and corresponding DCNN solvers are called inverse models. However, when the forward model of the system changes, the inverse model (i.e. the DCNN) has to be retrained. Moreover, for each forward model, a specialized inverse model needs to be trained.

To address the aforementioned shortcomings, we propose “*Physics Augmented U-Net*” which combines SIM and DCNNs as a generative prior. Our architecture involves three main elements: 1) modified U-Net model 2) a physics-based forward model of SIM and 3) an inference algorithm combining aforementioned two models. Here, the U-Net architecture is similar to the seminal work [24], but in the bottleneck two latent vectors are concatenated, as shown in Figure 2. The first latent vector is the feature representation of the low-frequency image (LFLV), learned by the encoder in the U-Net architecture. The second latent vector is used to inject high-frequency information (HFLV) acquired by the microscope to the network in order to generate high-resolution images. In the inference stage, high-frequency encoder is removed and HFLV is optimized by minimizing the loss between the measured image by the microscope and the output of the forward model for the generated image by the U-Net. Compared to traditional solvers, our approach is less computationally complex since we optimize a latent vector, instead of optimizing every pixel of the image. Our proposed approach is forward model agnostic to avoid retraining of the model. The proposed model is also more stable compared to other generative prior-based methods.

**Figure 1.**
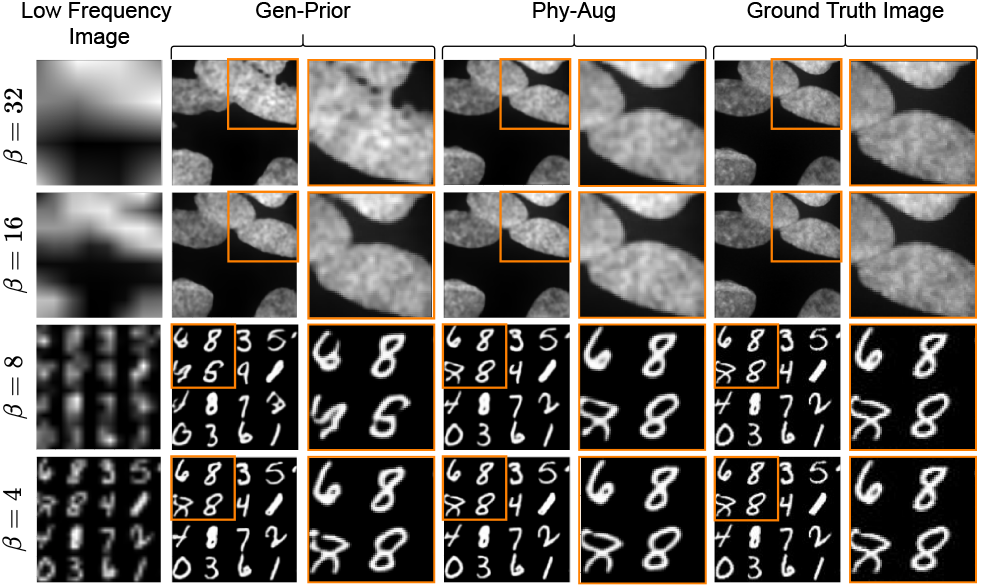
Visualization of the generated images by our proposed *Physics Augmented U-Net* on U2OS Cells and PatchMNIST datasets. Low-frequency image is generated from the ground truth patterned image stack. Gen-Prior is generated by the modified U-Net, given the input of low-frequency image and random noise vector for HFLV. Phy-Aug represents the final output image of our proposed method.

**Figure 2.**
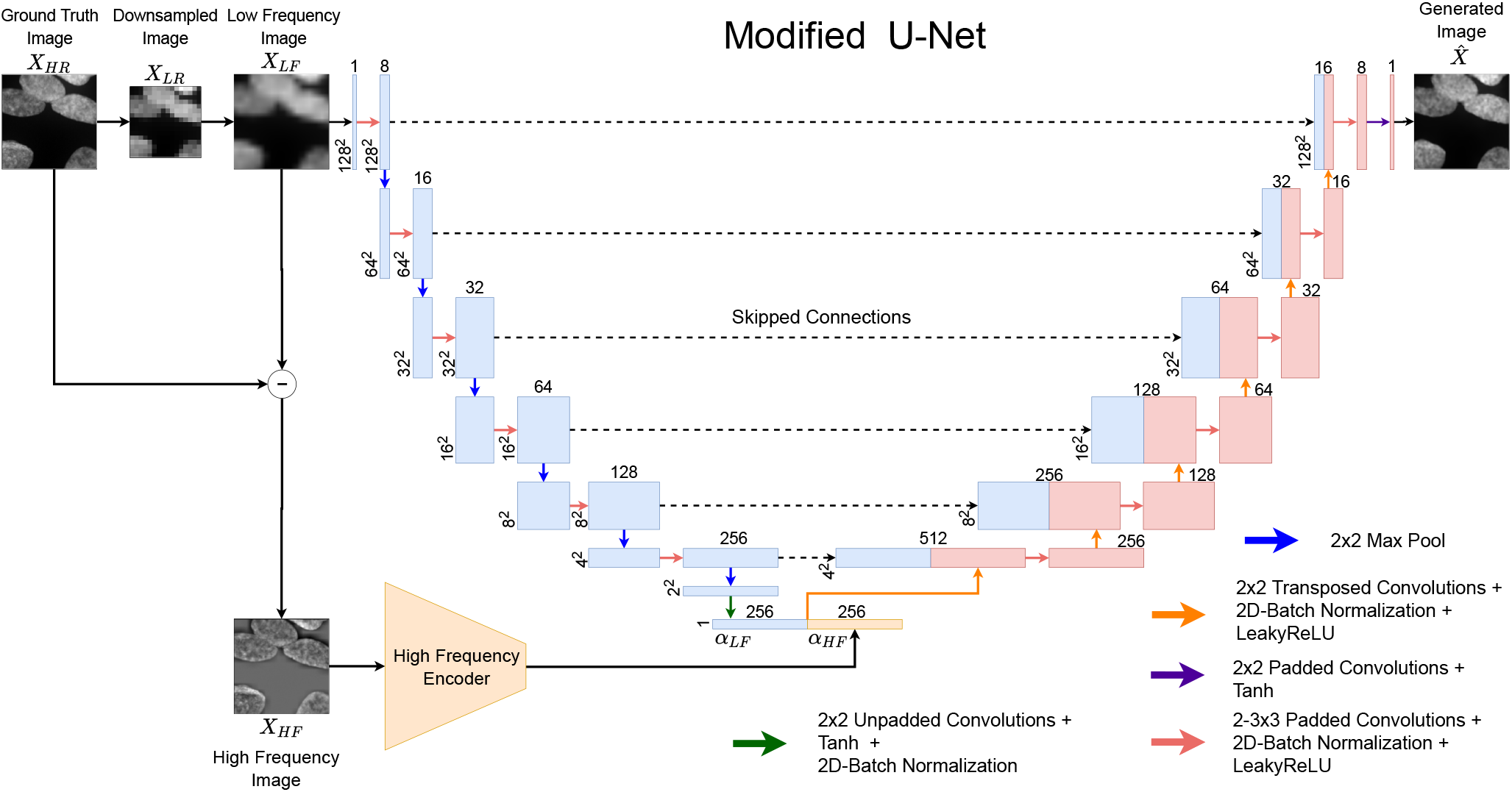
Architecture of the modified U-Net with high-frequency encoder during the training stage. Two latent vectors *α_LF_* and *α_HF_* are concatenated in the bottleneck of the modified-U-Net. The encodings for the low-frequency latent vector *α_LF_* is learned by the contracting path of the modified U-Net from *X_LF_* and the encodings for high-frequency latent vector *α_HF_* is learned by the high-frequency encoder.

**Our main contributions can be summarized as follows :**

- Proposing a novel method, termed *Physics Augmented U-Net* by combining deep learning and SIM for high-resolution microscopic image generation independent of the variation of SIM.
- High-frequency content aware modified U-Net architecture, with two latent vectors in the bottleneck - LFLV and HFLV, in which HFLV is optimized in the inference mode.
- Inference algorithm, to further improve the generated prior image by the modified U-Net by optimizing the HFLV to fit the measured microscopic images.

## 2. Related works

### Deep generative priors for super resolution^1^

Deep learning generative models are great at capturing the statistical information from large data as a prior. Such deep generative priors can be used in solving the inverse problems such as image reconstruction. A stronger prior will be able to reconstruct even a largely downsampled image into a super-resolved image. Standard generative models are designed to take a random noise vector as the input and output the generated image. Reusing such pre-trained models as priors is challenging. GAN inversion i.e., inverting the input image to a latent vector such that it can be reconstructed by the generative model, is the common practice to address this problem [4, 8, 16, 20]. GAN inversion falls into two categories: 1) optimizing the latent vector [6, 8, 15, 16, 20] 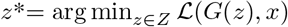, where *Z* denotes the latent vector space, *G* denotes the trained deep generative model, *x* corresponds to the low-resolution image and 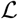 denotes the task-specific loss function. The super resolved image can be reconstructed from the optimal latent vector; 2) training an additional encoder to capture the mapping function from image space to latent vector space [1, 4, 21]. For instance, mGANprior [8] employs and optimizes multiple latent vectors to improve the reconstruction performance of the model. PULSE [16] iteratively optimizes the latent vector to generate high-resolution realistic images using StyleGAN. DGP [20] reduces the distance between training and testing image distributions by moderately fine-tuning the parameters of generator along with the latent vector. Meanwhile, GLEAN [4] applies an encoder to extract both the latent vectors and multi-resolution convolutional features, to learn significant high-level features and to provide additional spatial guidance for reconstruction. In contrast, Physics Augmented U-Net utilizes two latent vectors, where the first latent vector which provides low-frequency information is learned by the modified U-Net and the second latent vector which is used to inject high-frequency information is optimized by the inference algorithm. Our optimization process improves the high-frequency content in the reconstructed image and thus, the quality of resolution enhancement is improved.

### Deep Learning based SIM

In recent years, deep learning approaches have played a major role in biomedical image analysis. Microscopic imaging too has benefited from deep learning, with deep neural networks applied to super resolution [5, 11, 14, 18, 19, 23, 27, 29], image restoration [33], deconvolution [25,32], de-scattering [31], phenotyping [17], classification and image segmentation. Existing super resolution researches include application of deep learning for regular optical microscopes [23], PALM [19], STORM [18], Fourier ptychographic microscopy [27] and SIM [5,11,14]. Belthangady et al. [2] has conducted a comprehensive review on this line of work and presented the promises and pitfalls deep learning pose in the microscopic image reconstruction domain.

SIM acquires many raw images with different illumination patterns for high-resolution image generation. Therefore, recent researches on SIM have focused on reducing the number of raw images for image reconstruction, by which the speed of image acquisition can be increased and the number of times a specimen gets exposed can be reduced. Ling et al. [14] used cycle-consistent generative adversarial network (CycleGAN [37]) for image reconstruction. Jin et al. [11] used deep neural networks for super resolution image reconstruction from low light SIM images as well as from reduced number of raw images. Christensen et al. [5] used deep learning for image reconstruction from synthetic raw SIM images. To the best of our knowledge deep generative priors have not been used for image reconstruction in SIM.

### Image super resolution ^1^

Most of the super resolution algorithms [7,9,34,35] are based on single image super resolution as they learn the mapping between low-resolution and high-resolution images, and optimize the pixel-wise mean squared error (MSE). The resolution enhancement achievable through such super resolution algorithms are limited as they depend on the pixel-wise constraints which lead to perceptually unconvincing output with blurry effects on the edges due to their ill-posed nature [3, 13]. The generative adversarial network (GAN) based super resolution [13,22,30] alleviates the problem since they learn the mapping between the distributions of low-resolution and super resolution images, and approximates the natural image manifold, which results in realistic image generation. Recently, GAN-based image super resolution works focus on achieving large upsampling factors such as ×8, ×16 [10,26] instead of typical upsampling factors like ×2, ×4.

## 3. Physics Augmented U-Net

This section gives detailed introduction to our proposed method *Physics Augmented U-Net*. SIM is capable of capturing some high frequencies even when the image is sampled at low-resolution. In order to utilize the captured high-frequency information and to be forward model agnostic, our method consists of three elements: 1) modified U-Net architecture, 2) physics-based forward model of SIM and 3) inference algorithm.

### 3.1. Modified U-Net Architecture

Let’s formally define our low-resolution to high-resolution image translation task by the modified U-Net. Given high-resolution ground truth image 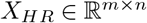 is down-sampled by average pooling operation to generate the corresponding low-resolution image 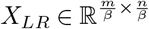, where *m, n* are the dimensions of the image and *β* denotes the downsampling factor.

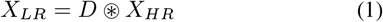

Here, *D* is the kernel used for average pooling and ⊛ denotes the convolution operation. Then, 2D bilinear upsampling operation with scaling factor of *β* is applied to *X_LR_* to generate the corresponding low-frequency image 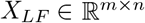. We generate a high-frequency image 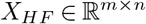 corresponding to *X_HR_* by subtracting *X_LF_* from *X_HR_*.

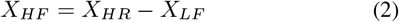

The input space to our modified U-Net consists of {*X_LF_, X_HF_*} pairs and the output space consists of *X_HR_*. The goal is to learn an image-to-image translation function *G_θ_*: {*X_LF_, X_HF_*} → *X_HR_* by minimizing the error rate: 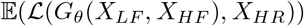. Here 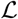 denotes the loss function, which is the mean absolute error (MAE) and *θ* represents the parameters of the modified U-Net, *G*. Let 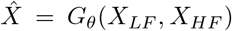 be the generated image by the U-Net, the loss function 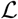:

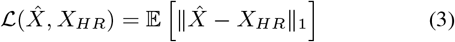

The architecture of the modified U-Net as shown in Figure 2, is similar to the seminal work [24], consists of the contracting path to learn the encodings of LFLV from input *X_LF_* and symmetric expanding path for high-resolution image 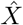 generation. We employ skipped connections between similar convolutional network blocks in the contracting and expanding paths. Additionally, a second encoding path is employed named *High-Frequency Encoder* with the similar structure of the contracting path to learn the encodings of HFLV from input *X_HF_*. LFLV and HFLV are concatenated in the bottleneck of the U-Net to enhance low-resolution to high-resolution image translation by injecting high-frequency information.

The contracting path consists of repeated convolutional network blocks with two 3×3 padded convolutions, each followed by 2D batch normalization and leaky rectified linear unit (LeakyReLU). After each convolutional network block a 2×2 max pooling operation is employed for downsampling. Six convolutional network blocks followed by max pooling operation are utilized to structure the contracting path. The number of feature channels are doubled at each downsampling step. Then an additional 2×2 unpadded convolution followed by tanh activation unit and batch normalization is applied to acquire LFLV. The tanh activation is applied to constraint the range of LFLV and then normalized using batch normalization to smoothen its vector space. The structure of encoding path in *High-Frequency Encoder* to acquire HFLV is same as the contracting path mentioned above without skip connections.

The expanding path consists of 2×2 transposed convolution operation for upsampling followed by 2D batch normalization and LeakyReLU. Then the upsampled feature map is concatenated with the corresponding feature map in the contracting path. Convolutional network block similar to contracting path is applied after each upsampling step and the number of feature channels were reduced by half. Six upsampling steps followed by convolutional network block are utilized to structure the expanding path. An additional 3×3 padded convolution operation followed by tanh activation is applied in the final layer to acquire the high-resolution image.

### 3.2. Physics-Based Forward Model

The task of the physics-based forward model is to simulate the corresponding SIM observation for a given high-resolution image generated by the modified U-Net. For demonstration in this paper, we use a SIM with random structured illuminations, similar to that used in DEEP-TFM [36]. We use a set of random binary patterns with the same size of the image as structured illuminations. The advantage of selecting random patterns is that they contain all frequencies [36]. Let *F_SIM_* be the physics based forward model. *F_SIM_* contains a randomized block of binary patterns 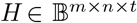, where 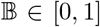; *t* represents the number of patterns. The randomized block of binary patterns are generated by creating matrices of size *m*×*n* and randomly assigning binary values to each of its pixels. Then the matrices are stacked together to get the randomized block of binary patterns. Finally, pixel-wise multiplication operation is employed between 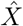 generated by the modified U-Net and each patterns in the randomized block of binary patterns *H* to acquire patterned image stack 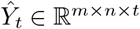.

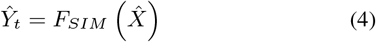

where,

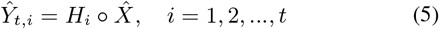

Here, 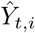 denotes the *i*th structured illuminated image before undersampled detection; ◦ denotes pixel-wise multiplication. Then, each structured illuminated image in 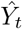 is downsampled by average pooling operation to generate 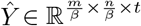. 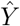 is the final simulated image corresponding to the physical observations.

### 3.3. Inference Algorithm

In the real application, we do not have access to the high-frequency image *X_HF_*. The high-frequency information required by the modified U-Net to generate high-resolution image should be injected externally. Our proposed inference algorithm as shown in Figure 3, focuses on utilizing the high frequencies captured by the SIM by combining both modified U-Net and physics-based forward model of SIM. In the inference algorithm, *High-Frequency Encoder* in the modified U-Net is removed. The low-frequency input image to the modified U-Net is generated using the measured image stack captured by SIM based microscope. Let 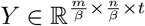 be the measured image stack from the SIM. We generate 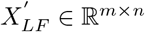 from *Y*. In order to have similar statistical distribution between 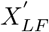 and *X_LF_* (training set), *Y* is first averaged along *t* axis to generate a 2D image 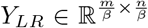 and then upsampled using 2D bilinear upsampling operation to create 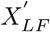 (similar to Section 3.1).

**Figure 3.**
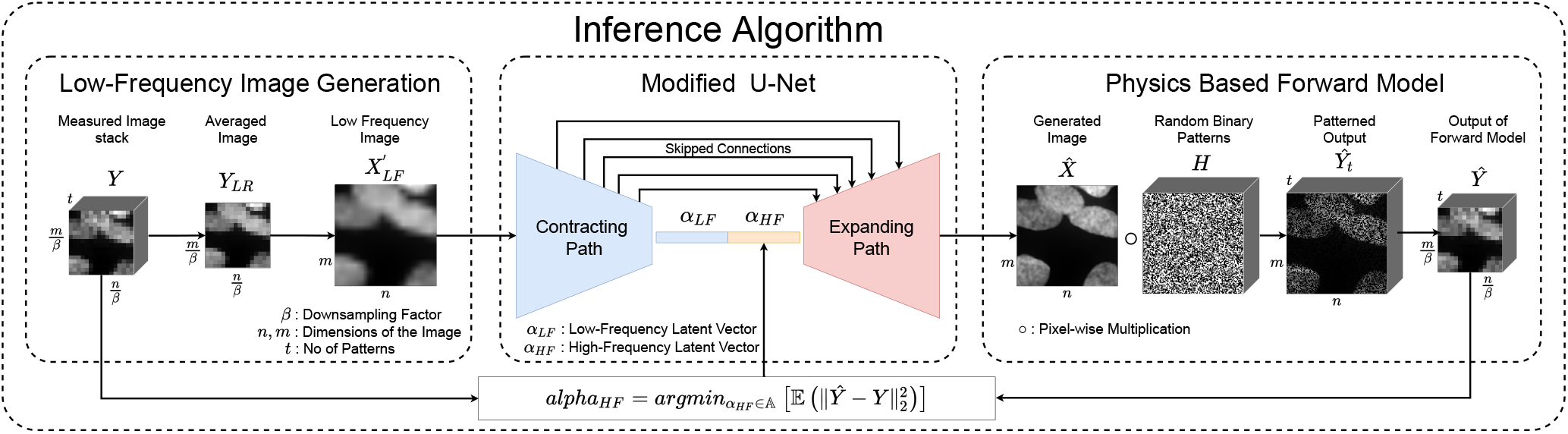
Illustration of our proposed *Physics Augmented U-Net* in the inference stage, consisting modified U-Net, physics-based forward model of SIM and inference algorithm to optimize *α_HF_* to obtain high-resolution image. The process of obtaining low-frequency input image is illustrated under low-frequency image generation block.

Let HFLV of the modified U-Net be 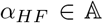, where 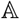 denotes the latent vector space. Initially, a prior image 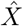 is acquired by feeding 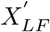 to the modified U-Net and randomly initializing *α_HF_* in the bottleneck. Here, random values for *α_HF_* is sampled from standard normal distribution 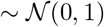. Then 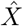 is fed to *F_SIM_* to acquire its corresponding image stack 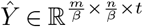, which is equivalent to SIM measurements. Iteratively, *α_HF_* is optimized by minimizing the loss 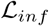 between *Y* and 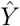. Here, MSE is employed as 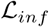. The goal is to inject high-frequency information captured by SIM to achieve high-resolution images by optimizing *α_HF_* such that:

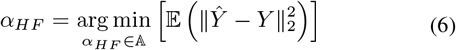

We employ Adam optimizer [12] with the learning rate of 0.1 during inference.

## 4. Experiments

We validated our method using two datasets: 1) PatchMNIST dataset and 2) U2OS Cells dataset, under different experimental setups. Details of the datasets and experimental setups used for validation are mentioned under Section 4.1 and 4.2.

### 4.1. Datasets

#### PatchMNIST Dataset

We generated a new dataset - PatchMNIST dataset, from the publicly available MNIST database^*^. MNIST database comprises of handwritten digits and it is commonly used for evaluating learning techniques and pattern recognition methods; it requires minimal efforts for data preparation. MNIST images are of size 28×28, but our architecture takes 128×128 images as the input. Since our model is used to generate high-resolution images, upsampling MNIST images to the required size is not appropriate; upsampled images will not have any high-frequency information. As a solution, we generated PatchMNIST dataset, which retains high-frequency information and poses a more challenging evaluation task. Under PatchMNIST data generation, MNIST images are resized to 32×32 and 400 such images are tiled together to form 20×20 image grids. From each of the resultant 640×640 image grids, 32 patches of size 128×128 are extracted to form the dataset. The generated dataset is then split into train, validation and test sets of 192000, 3200 and 100 images respectively. Train and validation sets are utilized for training and fine-tuning of the modified U-Net and high-frequency encoder. The test set is utilized to evaluate the performance of Physics Augmented U-Net.

#### U2OS (Bone Osteosarcoma) Cells Dataset

U2OS (bone osteosarcoma) cells were fixed with 4% paraformaldehyde and stained with DAPI. The cells were then imaged using a spinning disk confocal microscope at 63× magnification using an objective with 1.4 numerical aperture. Image stacks of size 60×2304×2304 were utilized for our performance evaluation. From each image stack, the corresponding maximum projection image is extracted and intensity transformation is applied. The resultant images are further downscaled by 63/20 and from each image 100 patches of size 128×128 are extracted to form the U2OS Cells dataset. The generated dataset is then split into train, validation and test sets of 14700, 4200 and 100 images respectively.

### 4.2. Experimental Setup

#### Training of Modified U-Net

The modified U-Net with the high-frequency encoder is trained end-to-end to perform low-resolution to high-resolution image translation task. The size of the latent vectors LFLV and HFLV is selected as 256. The model is trained using Adam optimizer [12] with learning rates of 1×10^−4^, and 2×10^−4^ for modified U-Net and high-frequency encoder respectively. *Beta1* and *beta2* of the Adam optimizer are set set to 0.9 and 0.999. The batch size is experimentally chosen as 32. We used MAE between the generated image and the ground truth as the loss function for training and the number of epochs is set to 500. The model is implemented in Pytorch ^†^ environment and trained using a Nvidia Tesla V100 PCIe graphics card with 32 GB memory.

The downsampling factor *β* used to create the low-frequency image *X_LF_* is varied to evaluate the capabilities of the proposed method to reconstruct high-resolution image. The high-frequency content in *X_LF_* is reduced by increasing *β*, so that the low-resolution to high-resolution image translation becomes challenging. To evaluate the performance of our proposed method with *β* values of 4, 8, 16 and 32, the modified U-net was trained with all said *β* values.

#### Inference Algorithm

During inference mode, the generated image is fed to the physics-based forward model and the loss between the resultant output image stack and the measured image stack from SIM is minimized. For experiments, the ground truth image is fed to the forward model and downsampled ground truth patterned image stack is generated as the equivalent of measured image stack from SIM. Then MSE loss between downsampled ground truth patterned image stack and forward model output of the generated image is calculated for optimization. In the real application, illumination patterns used for imaging are known, thus same patterns are used in the forward model for inference. To replicate this, in the experiments, same patterns are used in the forward model for both ground truth and the generated images. The number of patterns used in the forward model is varied to evaluate our method’s capacity in high-resolution image generation as well as to avoid the system becoming over determined. For every experiment during inference, different patterns are used to ensure that the proposed method is independent of the forward model.

## 5. Results and Discussion

We evaluate our proposed method on both U2OS Cells and PatchMNIST datasets. The qualitative results are illustrated in Figure 4. Quantitative results are illustrated using PSNR and SSIM metrics. PSNR measures the average pixel difference between ground truth and generated images; SSIM measures the structural similarity between ground truth and generated images. Under performance evaluation, downsampling factor (*β*) and the number of illumination patterns (*t*) are varied, and the aforementioned metrics are calculated between the ground truth image and the following images:

- **Low-Frequency Image** 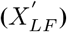, generated from the ground truth patterned image stack as mentioned under Section 3.3.
- **Generative Prior Image (Gen-Prior)**, generated by feeding 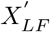 to the modified U-Net without the high-frequency encoder (here a random noise vector is fed as the HFLV).
- **Final Reconstructed Image (Phy-Aug)**, generated by feeding 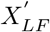 to our proposed Physics Augmented U-Net, comprising of modified U-Net, physics-based forward model and inference algorithm.

**Figure 4.**
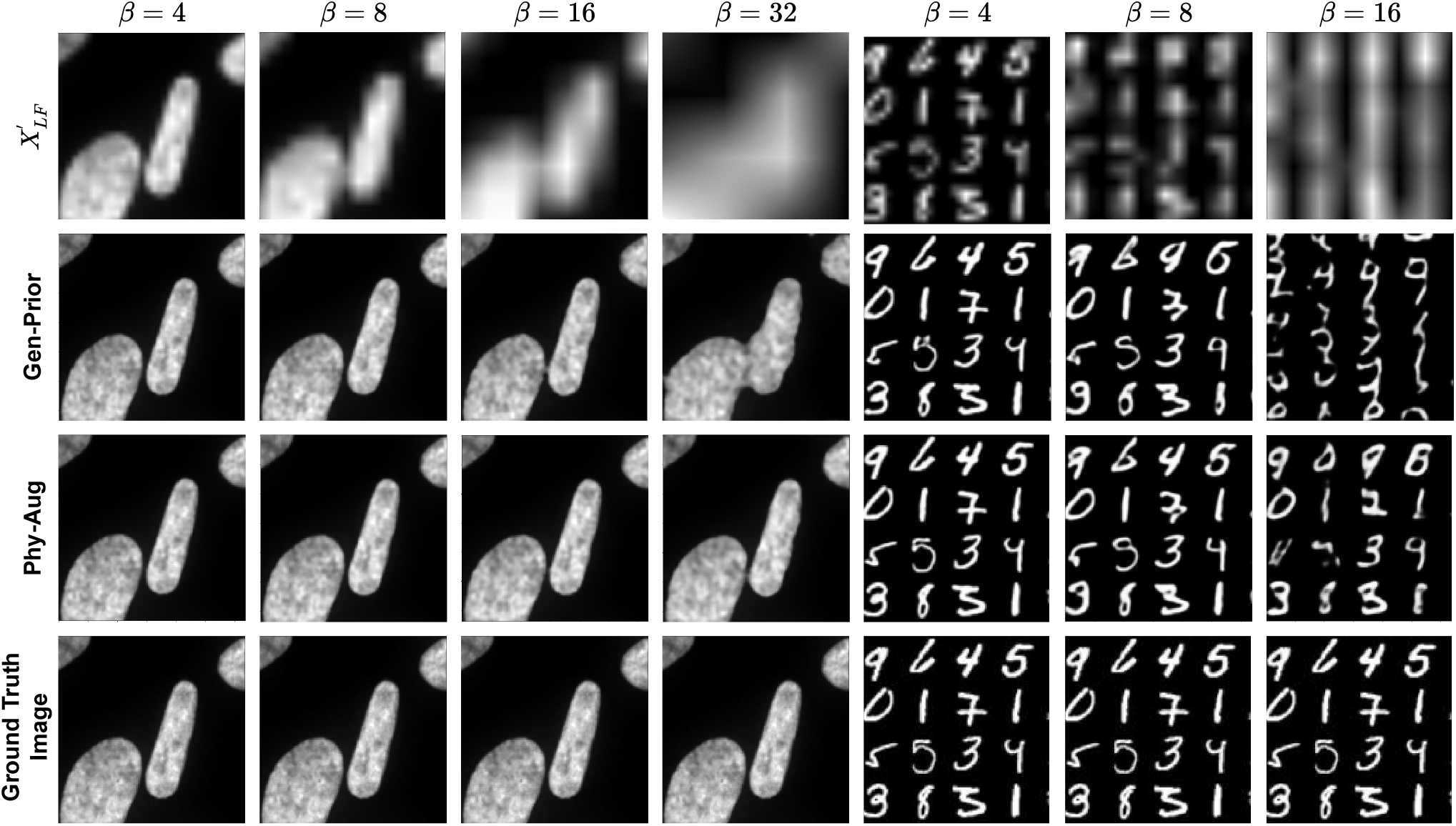
Illustration of the results on U2OS Cells and PatchMNIST datasets by our proposed *Physics Augmented U-Net* for different downsampling factors *β*. Number of illumination patterns (*t*) is set to 4. Significant improvement from the low-frequency input image 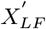 to Gen-Prior, and from Gen-Prior to Phy-Aug can be qualitatively observed. For additional results refer supplementary materials.

### 5.1. Performance with Different Downsampling Factors (*β*)

We investigated the performance of our proposed method for different *β* values. On the U2OS Cells dataset, we varied *β* by a factor of 2, as 4,8,16, and 32. Table 1 reports the performance of different *β* on the U2OS Cells dataset in terms of PSNR and SSIM metrics, when the number of SIM illumination patterns *t* = 4. From Table 1, it can be seen that Gen-Prior has significantly improved the low-frequency image 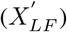 by an average improvement of 17.04 and 0.353 in PSNR and SSIM respectively, across the considered *β* values.

**Table 1.** Quantitative comparison (PSNR/SSIM) of the proposed method on **U2OS Cells dataset** for different downsampling factors (*β*), when number of patterns (*t*) is set to 4.

| Metrics | Methods | $\beta$ | | | |
| --- | --- | --- | --- | --- | --- |
|  |  | 4 | 8 | 16 | 32 |
| PSNR | $X'_{LF}$ | 26.55 | 21.15 | 16.29 | 10.24 |
|  | Gen-Prior | 40.75 | 36.05 | 35.29 | 30.31 |
|  | Phy-Aug | 42.22 | 39.58 | 37.42 | 36.48 |
| SSIM | $X'_{LF}$ | 0.876 | 0.738 | 0.555 | 0.269 |
|  | Gen-Prior | 0.989 | 0.973 | 0.968 | 0.921 |
|  | Phy-Aug | 0.992 | 0.987 | 0.979 | 0.973 |

This improvement by Gen-Prior is solely attributed to the modified U-Net architecture of our proposed method. Our proposed Physics Augmented U-Net was able to further enhance Gen-Prior by PSNR of 3.33 and SSIM of 0.020 on average. This improvement is attributed to the inclusion of physics-based forward model and inference algorithm for Phy-Aug generation.

On PatchMNIST dataset, we varied *β* by a factor of 2, from 4 to 16, i.e., *β* ∈ {4,8,16}. *β* = 32 is not considered for the experiments, because the size of each digit in a PatchMNIST image is 32×32 and downsampling it by a factor of 32 removes most spatial information. Table 2 reports the performance of different downsampling factors *β* on the PatchMNIST dataset when *t* = 4. From Table 2, it can be seen that Gen-Prior has significantly improved 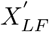 by an average improvement of 6.75 and 0.418 in PSNR and SSIM respectively, which was further enhanced by the Physics Augmented U-Net by PSNR of 1.64 and SSIM of 0.032 on average. Even with the downsampling factor of 16, a drop in performance can be seen from Table 2, which can be attributed to the loss of spatial information.

**Table 2.** Quantitative comparison (PSNR/SSIM) of the proposed method on **PatchMNIST dataset** for different downsampling factors (*β*), when number of patterns (*t*) is set to 4.

| Metrics | Methods | $\beta$ | | |
| --- | --- | --- | --- | --- |
|  |  | 4 | 8 | 16 |
| PSNR | $X'_{LF}$ | 14.68 | 10.24 | 3.56 |
|  | Gen-Prior | 23.57 | 15.69 | 9.47 |
|  | Phy-Aug | 26.72 | 17.49 | 9.44 |
| SSIM | $X'_{LF}$ | 0.565 | 0.194 | 0.026 |
|  | Gen-Prior | 0.916 | 0.761 | 0.362 |
|  | Phy-Aug | 0.934 | 0.818 | 0.384 |

### 5.2. Performance with Different No. of Illumination Patterns (*t*)

We also investigated the performance of our proposed method with different values for *t*. On both U2OS Cells and PatchMNIST datasets, we varied *t* from 2 to 32 by a factor of 2, i.e., *t* ∈ {2,4,8,16,32}. Table 3 reports the performance of different t on U2OS Cells dataset when *β* = 32 and on PatchMNIST dataset when *β* = 8. We observed increase in performance from our proposed Physics Augmented U-Net, with the increase of *t*. However, we were able to achieve good performance even with low number of patterns (*t* = 2) and thus image acquisition time of the microscope can be reduced. The forward model is independent of the modified U-Net, and thus our method gives similar results to any variation of random binary patterns used in the forward model without further retraining.

**Table 3.** Quantitative comparison (PSNR/SSIM) of the proposed method on both **U2OS Cells and PatchMNIST dataset** for different number of patterns (*t*).

| Dataset | $\beta$ | $t$ | Metrics | |
| --- | --- | --- | --- | --- |
|  |  |  | PSNR | SSIM |
| U2OS Cells | 32 | 32 | 38.171 | 0.9803 |
|  |  | 16 | 37.605 | 0.9777 |
|  |  | 8 | 36.820 | 0.9735 |
|  |  | 4 | 36.485 | 0.9726 |
|  |  | 2 | 36.491 | 0.9729 |
| PatchMNIST | 8 | 32 | 19.244 | 0.8524 |
|  |  | 16 | 18.783 | 0.8450 |
|  |  | 8 | 18.220 | 0.8334 |
|  |  | 4 | 17.491 | 0.8177 |
|  |  | 2 | 16.848 | 0.8024 |

### 5.3. Comparison with Traditional Optimization

We investigated the performance of Phy-Aug with the output image generated from the traditional optimization method used for resolution enhancement (Trad-Opt). For Trad-Opt generation, a 2D image 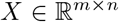 is randomly initialized and fed to the forward model, to obtain the corresponding image stack equivalent to SIM measurements. Then, every pixel of the 2D image *X* is optimized to fit the downsampled ground truth patterned image stack to the output of forward model as given below:

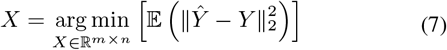

Adam optimizer is used for this experiment and *β, t* and number of iterations are set to 32, 2048 and 100000 respectively. *β, t* and number of iterations of Phy-Aug generation are set to 32, 4 and 2000 respectively. Figure 5 illustrates the comparison between Trad-opt and Phy-Aug along with the PSNR values. The results clearly state that Physics Augmented U-Net is capable of generating high resolution image faster than the traditional optimization method and requires lesser number of illumination patterns.

**Figure 5.**
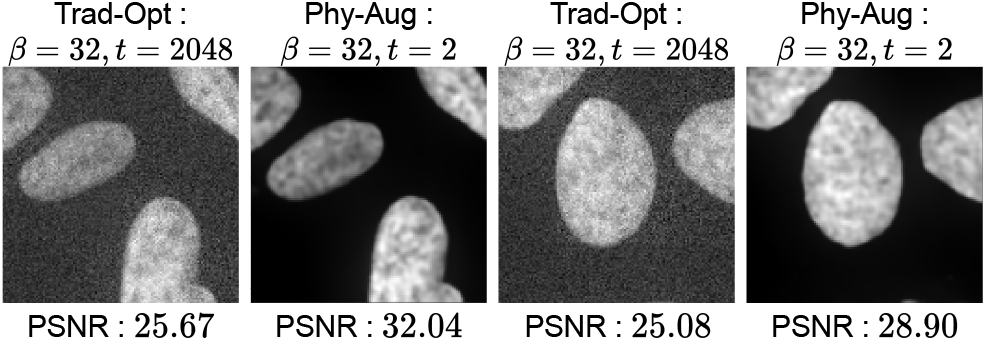
Comparison between Trad-Opt and Phy-Aug

### 5.4. Comparison with Modified U-Net with High-Frequency Encoder

We compared Phy-Aug with the output image from the combined architecture of modified U-Net and high-frequency encoder (M.UNet+HFE) and the corresponding quantitative results are provided in Table 4. M.UNet+HFE is the best possible high resolution image generated by feeding all ground-truth high-frequency information and thus, provides the best results. From Table 4, we can observe that our proposed method converges towards the best possible results.

**Table 4.** Quantitative comparison (PSNR) of Phy-Aug with M.UNet+HFE on **U2OS Cells dataset** for different downsampling factors (*β*), when number of patterns (*t*) is set to 4.

| Metrics | Methods | $\beta$ | | | |
| --- | --- | --- | --- | --- | --- |
|  |  | 4 | 8 | 16 | 32 |
| PSNR | M.UNet+HFE | 42.40 | 40.71 | 39.95 | 39.58 |
|  | Phy-Aug | 42.22 | 39.58 | 37.42 | 36.48 |

## 6. Limitations

Our modified U-Net is trained using low-frequency images. In order to utilize the trained U-Net during inference, the input image i.e., low frequency image obtained from SIM measurements should be statistically similar to the training image distribution. The SIM illumination patterns used for measurements should have the ability to generate such low frequency images, thus the variations of SIM patterns that can be used are limited.

Next, our method relies on the assumption that the prior generated from the modified U-Net accurately corresponds to the input low-frequency image. However, for some cases, we observed good performance with priors not being accurate estimates of the input low-frequency images. But, we have not carried out an extensive sensitivity analysis to quantify that and thus our method is limited by this constraint.

## 7. Conclusions and Future Work

In this paper we propose “Physics Augmented U-Net”, to enhance the resolution of undersampled structured illumination microscopy (SIM) images. Importantly, rather than learning an image translation from under-sampled low-resolution image space to the high-resolution image space, we train a U-Net-like architecture as a generative model. A latent vector corresponding to high-resolution information can then be further optimized to infer high-resolution images from the prior, through a physics-based forward model of SIM. In contrast to traditional SIM solvers that optimize over the entire 2D image, Physics Augmented U-Net optimizes a much smaller latent vector, thus resulting in lower computational complexity. Since the forward model is explicitly separated from the learned prior, Physics Augmented U-Net also works with any variation of SIM without further retraining. Compared to other adversarially-trained generative models, the proposed approach is much stable. In future work, we will implement more realistic forward models as well as extend to different SIM variations and datasets. Given its similarity to the seminal U-Net architecture, we anticipate that practitioners of microscopy can easily adapt Physics Augmented U-Net for new datasets as well as for any variation of SIM.

## Supplementary Material

### 1. Qualitative Comparison Between Low Frequency Image, Gen-Prior, Phy-Aug and Ground Truth Images

. We investigated the performance of Physics Augmented U-Net on U2OS Cells dataset and the corresponding qualitative results are illustrated in Figure 6. Even with an under-determined system with *β* = 32 and *t* =4, our proposed architecture was able to yield good performance. Our modified U-Net itself was able to generate Gen-Priors with significant qualitative improvement and the inference algorithm further improved the quality by incorporating the optimal high-frequency information. The high frequency information of the image being captured efficiently is qualitatively illustrated in Figure 6.

**Figure 6.**
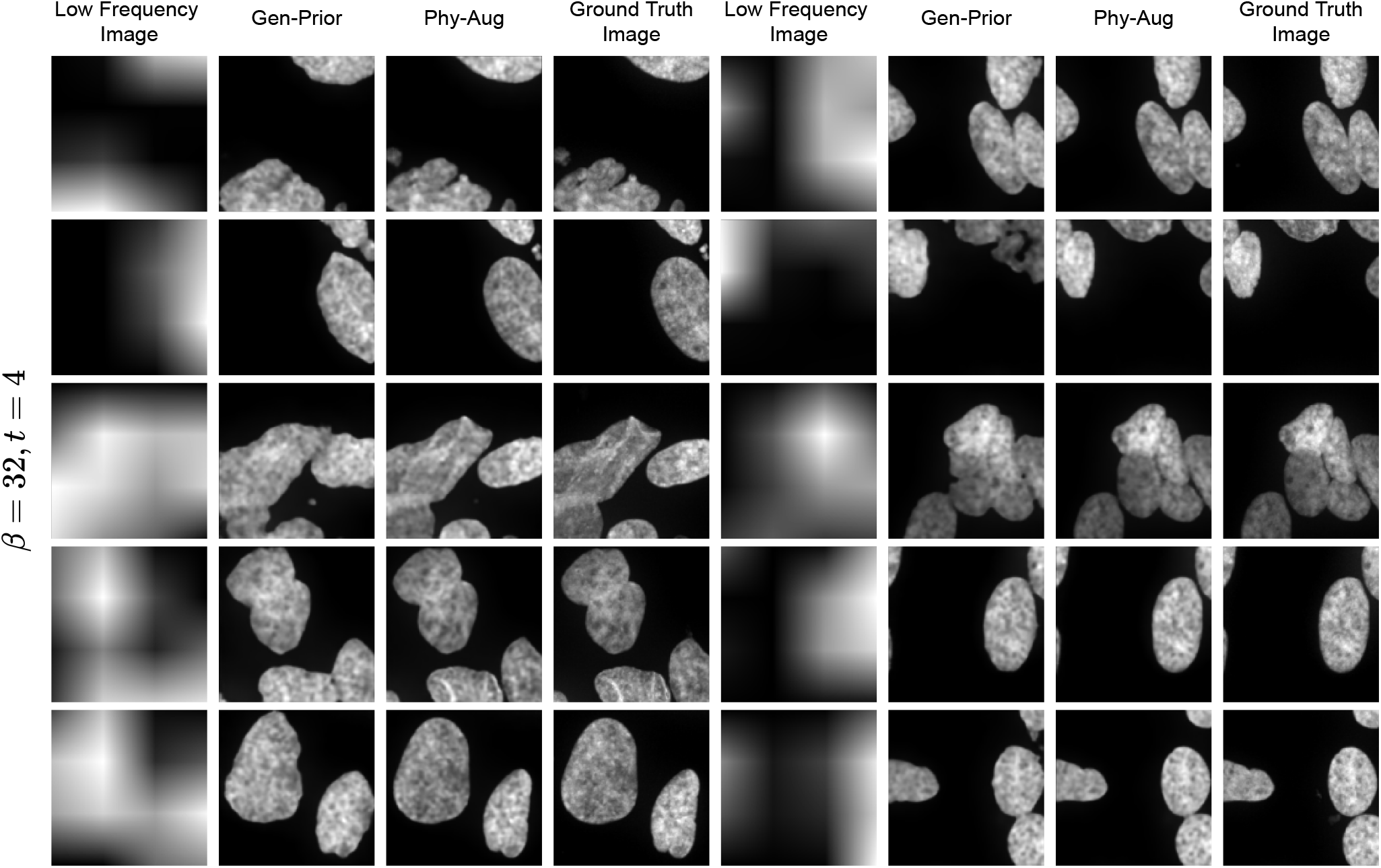
Illustration of our proposed architecture - *Physics Augmented U-Net’s* performance on U2OS Cells dataset for downsampling factor (*β*) of 32 and number of illumination patterns (*t*) of 4. Significant improvement from the low-frequency input image 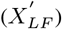 to Gen-Prior, and from Gen-Prior to Phy-Aug can be qualitatively observed.

### 2. Performance Analysis with Different Downsampling Factors (*β*) and No of Illumination Patterns (*t*)

. We compared the performance of the final reconstructed image (Phy-Aug) with the output image from the combined architecture of modified U-Net and high-frequency encoder (M.UNet+HFE) and Gen-Prior under different *β* and *t* values. The corresponding quantitative results are illustrated in Figure 7 and 8, using peak signal-to-noise ratio (PSNR) and structural similarity index (SSIM) metrics. As t is increased, we observed increase in performance from our proposed method as well as convergence towards M.UNet+HFE, which is the best possible high resolution image generated by feeding all the high frequency information. This indicates that our method comprising of modified U-Net, physics based forward model and inference algorithm is capable of incorporating the optimal high-frequency information for resolution enhancement of a low frequency input image.

**Figure 7.**
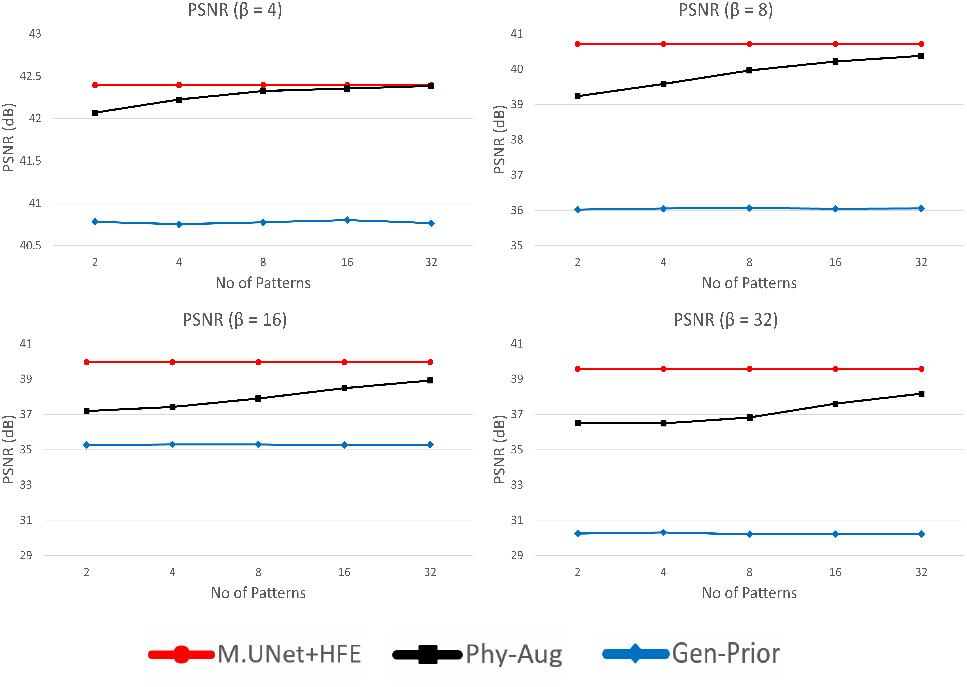
Quantitative comparison (PSNR) of Phy-Aug with Gen-Prior and M.UNet+HFE on **U2OS Cells dataset** for different number of patterns (*t*) and downsampling factors (*β*). For better illustration, y-axis range of the plot is varied.

**Figure 8.**
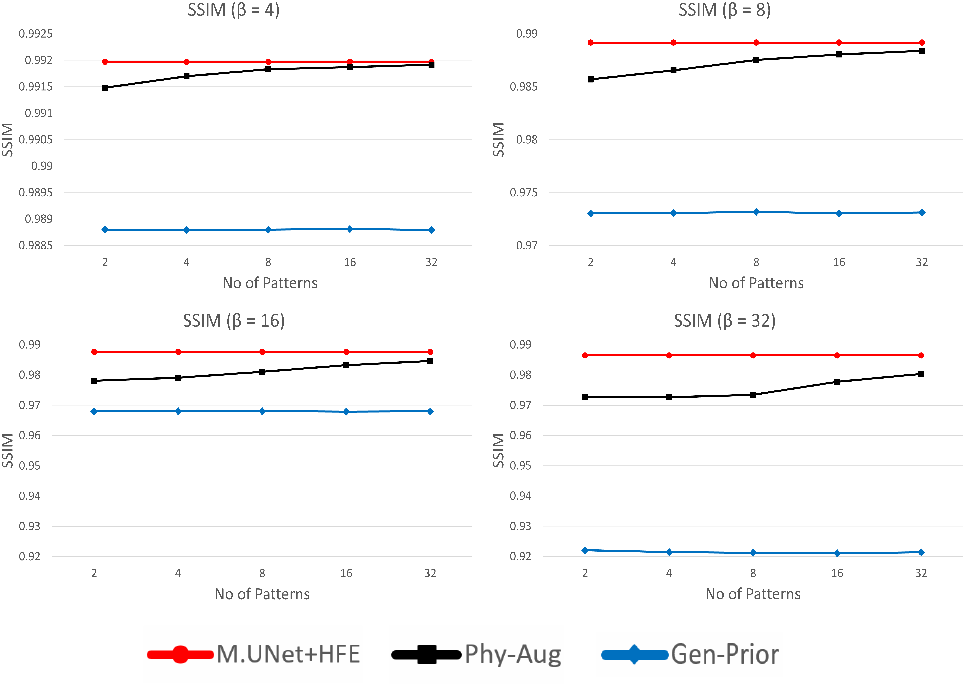
Quantitative comparison (SSIM) of Phy-Aug with Gen-Prior and M.UNet+HFE on **U2OS Cells dataset** for different number of patterns (*t*) and downsampling factors (*β*). For better illustration, y-axis range of the plot is varied.

## Footnotes

1 Here the term *super resolution* refers to high-resolution images reconstructed from low resolution measurements. We do not refer to superresolution microscopy that can image beyond the diffraction limit.

1 As stated before, here the term *super resolution* refers to high-resolution images reconstructed from low resolution measurements, not images beyond the diffraction limit.

* http://yann.lecun.com/exdb/mnist/

† https://pytorch.org/

## Notes

### Competing Interest Statement

The authors have declared no competing interest.

